# Familiarity increases processing speed in the visual system

**DOI:** 10.1101/670489

**Authors:** Mariya E. Manahova, Eelke Spaak, Floris P. de Lange

## Abstract

Familiarity with a stimulus leads to an attenuated neural response to the stimulus. Alongside this attenuation, recent studies have also observed a truncation of stimulus-evoked activity for familiar visual input. One proposed function of this truncation is to rapidly put neurons in a state of readiness to respond to new input. Here, we examined this hypothesis by presenting human participants with target stimuli that were embedded in rapid streams of familiar or novel distractor stimuli at different speeds of presentation, while recording brain activity using magnetoencephalography (MEG) and measuring behavioral performance. We investigated the temporal and spatial dynamics of signal truncation and whether this phenomenon bears relationship to participants’ ability to categorize target items within a visual stream. Behaviorally, target categorization performance was markedly better when the target was embedded within familiar distractors, and this benefit became more pronounced with increasing speed of presentation. Familiar distractors showed a truncation of neural activity in the visual system, and this truncation was strongest for the fastest presentation speeds. Moreover, neural processing of the target was stronger when it was preceded by familiar distractors. Taken together, these findings suggest that truncation of neural responses for familiar items may result in stronger processing of relevant target information, resulting in superior perceptual performance.

**Significance statement:** The visual response to familiar input is attenuated more rapidly than for novel input. Here we find that this truncation of the neural response for familiar input is strongest for very fast image presentations. We also find a tentative function for this truncation: the neural response to a target image that is embedded within distractors is much greater when the distractors are familiar than when they are novel. Similarly, target categorization performance is much better when the target is embedded within familiar distractors, and this advantage is most obvious for very fast image presentations. This suggests that neural truncation helps to rapidly put neurons in a state of readiness to respond to new input.

## Introduction

The brain rapidly processes an immense amount of sensory information and is constantly engaged in learning, adapting its responses based on new knowledge of the environment. A large body of evidence (e.g., Fahy et al., 1993; Li et al., 1993; Sobotka and Ringo, 1993; Xiang and Brown, 1998; Freedman et al., 2006; Mruczek and Sheinberg, 2007; Anderson et al., 2008; Woloszyn and Sheinberg, 2012; Huang et al., 2018) has demonstrated that familiarity with a stimulus modulates neural processing in various ways.

One recently described consequence of familiarity is temporal sharpening, or ‘truncation’, of the signal evoked by a familiar stimulus (Meyer et al., 2014). When the visual system is familiar with a stimulus, it is able to process the stimulus more quickly, leading to a truncated response for familiar compared to novel images. Put differently, activity in the visual system returns to baseline levels more rapidly for familiar than for novel images, thus leaving the visual system more aptly poised to process future input. This may lead to faster, more efficient processing of new information, and a behavioral consequence may be that familiar images have lower saliency than novel ones. Indeed, monkeys as well as humans spend less time looking at familiar than novel images (Jutras and Buffalo, 2010; Ghazizadeh et al., 2016), and familiar distractors interfere with behavioral tasks more than novel ones (Mruczek and Sheinberg, 2007).

When images are presented in rapid succession, image truncation leads to a larger dynamic range (i.e., baseline-to-peak difference) for a stream of familiar than novel images (Meyer et al., 2014). In such a case, neural processing has not yet terminated for one novel image when the next one is presented, effectively reducing the ability of the visual system to respond to new input. In a previous MEG study in humans (Manahova et al., 2018), we observed a larger dynamic range in the visual response to familiar object items compared to novel ones localized in the lateral occipital complex (LOC), thus demonstrating that signal truncation also occurs in the human brain.

While signal truncation appears a robust and potentially useful phenomenon, there are still several outstanding questions. First, it is not clear to what extent signal truncation is present at different stages in the visual system. It may be a mechanism that underlies familiarity in the whole visual system, or it could be specific to object-selective cortex such as human area LOC and macaque area IT (Meyer et al., 2014; Manahova et al., 2018). Second, the process’s temporal boundary conditions are unclear. Signal truncation has been observed for streams where stimuli were presented at a rate of 120-180 ms, but it is unknown whether the phenomenon extends to other temporal scales. Multiple studies show that processing within earlier cortical regions takes place at a faster intrinsic timescale than for later regions (Murray et al., 2014). Therefore, short image durations may potentially elicit the strongest signal truncation in early visual regions, while long image durations may do so in downstream visual areas. Third, if signal truncation aids in faster neural processing, then this phenomenon may be linked to behavioral improvements such as improved perceptual speed and accuracy.

To address these open questions, we examined the neural dynamics of stimulus processing along the visual hierarchy and their relationship to behavior. We embedded target stimuli in a stream of familiar or novel distractors while measuring human neural activity using MEG. To preview our findings, we found robust truncation of neural activity for familiar input, which was localized to early visual cortex for the shortest image duration and to later visual areas for medium-length image durations. Moreover, familiarity with the distractor stream led to stronger neural processing of the target and to superior behavioral performance, which was most pronounced for rapid visual streams. Together, our results suggest that neural truncation helps to rapidly put the visual system in a state of readiness to respond to new input.

## Materials and Methods

### Data and software availability

Data and code used for stimulus presentation and analysis will be made available online at the Donders Institute for Brain, Cognition, and Behavior repository. Videos displaying how different trial types appear to a participant will also be available there.

### Participants

Thirty-seven healthy human volunteers (25 female, 11 male, mean age = 26.4 years, *SD* = 6.6 years) with normal or corrected-to-normal vision, recruited from the university’s participant pool, completed the experiment and received monetary compensation. The sample size, which was defined a priori, ensured at least 80% power to detect within-subject experimental effects with an effect size of Cohen’s d>0.5. The study was approved by the local ethics committee (CMO Arnhem-Nijmegen, Radboud University Medical Center) under the general ethics approval (“Imaging Human Cognition”, CMO 2014/288), and the experiment was conducted in compliance with these guidelines. Written informed consent was obtained from each individual.

### Stimuli

Stimuli were chosen from the image sets provided at http://cvcl.mit.edu/MM/uniqueObjects.html and http://cvcl.mit.edu/MM/objectCategories.html. A different object was represented in each image, and all objects were shown against a gray background. A total of 554 images were used (20 target images, 6 familiar distractors, and 528 novel distractors). Familiar images were randomly selected for each participant and were shown during the behavioral training session as well as during the MEG testing session, while novel images were only shown during the MEG testing session. In both the behavioral and MEG sessions, the images subtended four by four degrees of visual angle.

### Apparatus

MATLAB (The Mathworks, Inc., Natick, Massachusetts, United States) and the Psychophysics Toolbox extensions (Brainard, 1997) were used to show the stimuli on a monitor with a resolution of 1920×1080 pixels and a refresh rate of 120 Hz for the behavioral session. For the MEG session, a PROpixx projector (VPixx Technologies, Saint-Bruno, QC Canada) was used to project the images on the screen, with a resolution of 1920×1080 and a refresh rate of 120 Hz.

### Trial structure

During the behavioral session as well as the MEG session, each trial began with a fixation dot (see Figure 1A for the trial structure). The fixation dot was presented for a randomly selected period between 500 and 750 ms. Then, a stream of images was presented, lasting for 2400 ms. The duration of the trial was kept the same, while the number of images per trial varied depending on the image duration. For images lasting 50 ms, there were 48 images per trial; for images lasting 100 ms, there were 24 images per trial; for images lasting 150 ms, there were 16 images per trial; for images lasting 200 ms, there were 12 images per trial; for images lasting 300 ms, there were 8 images per trial. At the end of the trial, if a correct response had been given (i.e., the target image had been categorized correctly as animate or inanimate), the fixation dot turned green for 500 ms. If the response was incorrect, the fixation dot turned red for 500 ms. Afterwards, a blank screen was presented for 1250 ms, and participants were encouraged to blink during this period.

**Figure 1.**
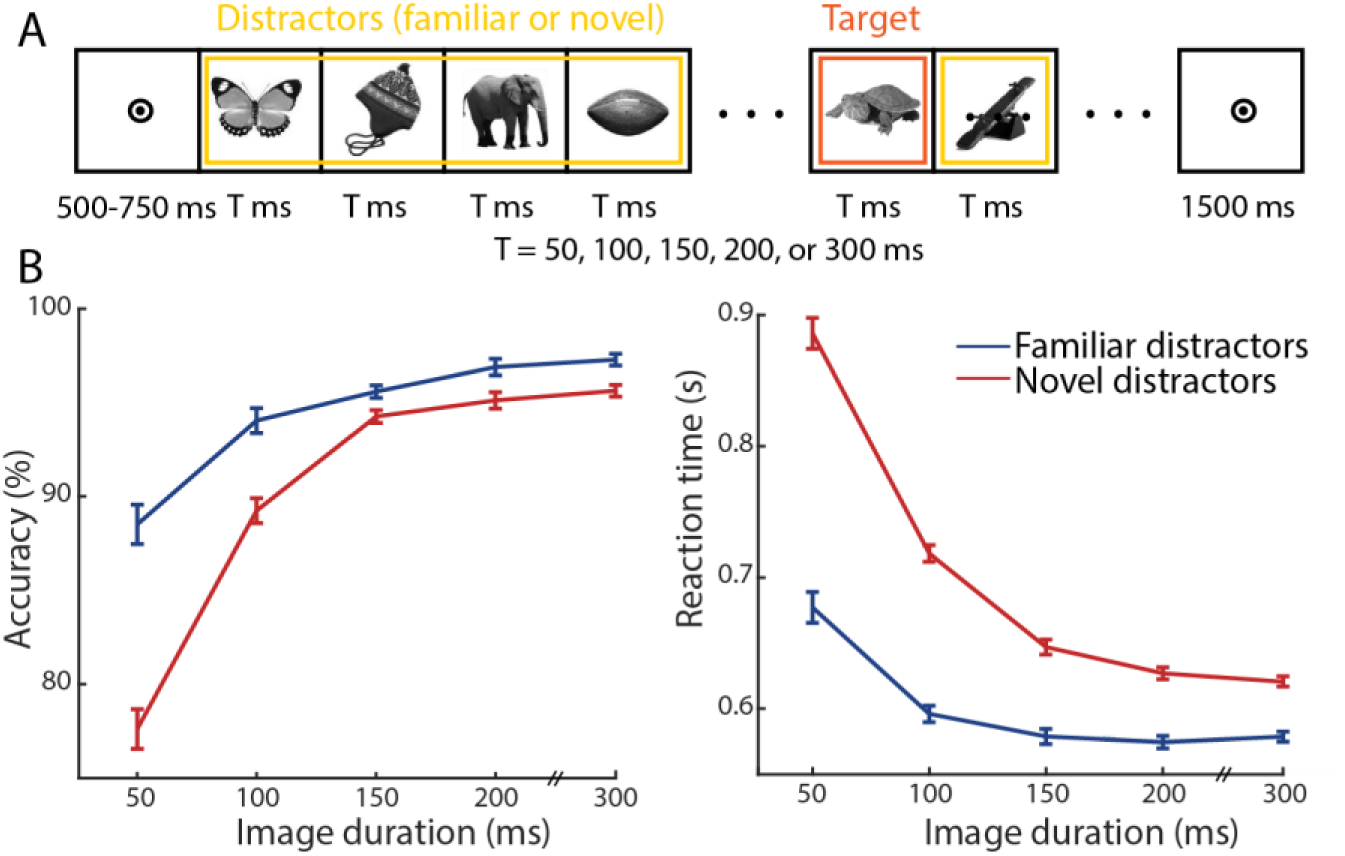
(A) Trial structure. First, participants saw a bull’s eye fixation point for a jittered period between 500 and 750 ms. Then, a stream of distractors was presented (here, highlighted in yellow), each image lasting for the allotted amount of time for that trial (50, 100, 150, 200, or 300 ms) and no gaps between images. Somewhere in the distractor stream, a target image was embedded (here, highlighted in orange). Participants had to respond as quickly as possible after detecting the target, categorizing the target image as animate or inanimate. Finally, a fixation dot was presented at the end of the trial, and participants received feedback on their performance. (B) Accuracy and reaction times per image duration. Blue lines denote familiar distractors, and red lines denote novel distractors. Error bars show the within-participant SEM.

### Experimental design

In the behavioral training session, participants were first taught to categorize the target images into animate or inanimate. There were twenty target images, and they saw them one at a time. The images were presented on the screen until a response was made. The participants pressed the left and right arrow keys to indicate their answer, with the response mapping randomized across participants. There were six blocks, each block showing all twenty target images. Participants were given feedback after each image about whether they categorized that image correctly. They were also shown their average accuracy at the end of each block.

Then, participants proceeded to practice the experimental task (see Figure 1A). They observed streams of images of distractors, and they had to detect the target image, which was one of the twenty images they had just learned. Once they detected the target among the distractors, they had to immediately press a button to categorize it as animate or inanimate. The target could occur at different time points in the trial. The trial lasted for 2400 ms, and on 90% of trials the target was presented sometime between 1200 ms and 2100 ms. On 10% of trials the target was presented before the 1200 ms mark of the trial to make sure that participants were paying attention throughout the trial and not only during the second half. These trials were discarded from further analysis. Notably, during the behavioral training session, the distractors were always the same six images. Since participants saw these repeatedly on every trial, they became highly familiar. Participants completed five blocks of 100 trials each for a total of 500 trials.

One or two days later, participants completed the MEG testing session in which they saw familiar (i.e., those presented during the behavioral session) and novel (not seen before) images. Half the trials in the experiment were familiar, and the other half were novel. On each trial, six images were chosen as distractors and repeated to match the number of images needed for that image duration (see Trial structure). If the trial was familiar, then these six images were the six distractors participants had seen repeatedly during the behavioral training session. If the trial was novel, the six images were randomly drawn (without replacement) from the set of novel images. Each novel image was shown four times during the course of the experiment; in contrast, each familiar image was displayed 350 times during the MEG session and 250 times during the behavioral session. The task was the same as during the behavioral training session: detect the target and categorize it as animate or inanimate immediately after seeing it. Participants completed seven blocks of 100 trials each for a total of 700 trials. At the end of the MEG testing session, participants’ knowledge of the familiar images was assessed. Participants saw 50 images, the six familiar ones and 44 selected at random from the novel images participants had been shown, and participants had to indicate whether the image was familiar or novel. ‘Familiar’ referred to images seen repeatedly during the behavioral training session as well as during the MEG testing session, while ‘novel’ referred to images seen only during the MEG testing session.

### Data acquisition

#### MEG recordings

Brain activity was recorded using a 275-channel MEG system with axial gradiometers (VSM/CTF Systems, Coquitlam, BC, Canada) in a magnetically shielded room. During the experiment, head position was monitored online and corrected if necessary (Stolk et al., 2013). Head position monitoring was done using three coils: one placed on the nasion, one in an earplug in the left ear, and one in an earplug in the right ear. MEG signals were sampled at 1200 Hz. A projector outside the magnetically shielded room projected the visual stimuli onto a screen in front of the participant via mirrors. Participants gave their behavioral responses via an MEG-compatible button box. Participants’ eye movements and blinks were also monitored using an eye-tracker system (EyeLink, SR Research Ltd., Mississauga, Ontario, Canada).

#### MRI Recordings

To allow for source reconstruction, anatomical magnetic resonance imaging (MRI) scans were acquired using a 3T MRI system (Siemens, Erlangen, Germany) and a T1-weighted MP-RAGE sequence with a GRAPPA acceleration factor of 2 (TR = 2300 ms, TE = 3.03 ms, voxel size 1 × 1 × 1 mm, 192 transversal slices, 8° flip angle).

### Data analysis

#### Preprocessing of MEG data

The MEG data were preprocessed offline using the FieldTrip software (Oostenveld et al., 2011). Trials where the target was presented earlier than 1200 ms into the trial (10% of trials) were removed from analysis. The data were demeaned, and noise was removed based on third-level gradiometers. Then, trials with high variance were manually inspected and removed if they contained excessive and irregular artifacts. This resulted in retaining, on average, 93% of trials per participant (range 86-99%). Afterwards, independent component analysis (ICA) was applied to identify regular artifacts such as heartbeat and eye blinks and remove the respective components.

#### Event-related fields

The data were filtered using a 6^th^ order Butterworth low-pass filter with a cutoff frequency of 40 Hz. Before calculating event-related fields (ERFs), the data were baseline-corrected on the interval starting at 200 ms before stimulus onset until stimulus onset (0 ms), i.e., the onset of the first distractor. For the ERF analysis of target-related data, the data were baseline-corrected on the interval starting at 100 ms before target onset until target onset (0 ms). Subsequently, the data were split into conditions based on the familiarity and the image duration for the trial. Then, the data were transformed to planar gradiometer representation in order to facilitate interpretation as well as averaging over participants. The familiar and novel conditions had an equal number of trials by design. The resulting ERFs were averaged over participants for visualization purposes. Standard error of the mean (SEM) was computed within participants (Cousineau, 2005) and with bias correction (Morey, 2008).

#### Power analysis

In order to assess the dynamic range (i.e., the baseline-to-trough difference) of the signal, we computed the power in the ERF as in (Manahova et al., 2018). First, we averaged the data over trials, timelocking to stimulus onset. The time window of interest was from 200 until 1200 ms. Next, we applied the planar transformation to the data. Then, we conducted a spectral analysis for all frequencies between 1 and 30 Hz with a step size of 1 Hz. We applied the fast Fourier transform to the planar-transformed time domain data, after tapering with a Hanning window. The power analysis was carried out separately per condition, where a condition was defined by familiarity and image duration. Afterwards, the horizontal and vertical components of the planar gradient were combined by summing. The resulting power per frequency was averaged over participants.

#### Source reconstruction

For maximal sensitivity, we carried out source reconstruction analysis. We used each participant’s anatomical MRI scan to create a volume conduction model based on a single-shell model of the inner surface of the skull (Nolte and Dassios, 2005). We computed subject-specific dipole grids, which were based on a regularly spaced 6-mm grid in normalized MNI (Montreal Neurological Institute) space. Then, the sensor-level axial gradiometer data were split into conditions determined by familiarity and image duration. For each condition, we carried out source analysis with the DICS method (Gross et al., 2001), quantifying coherence with a synthetic stimulus signal at the frequency of interest. We used coherence as a measure of dynamic range and thus signal truncation. Finally, we averaged the data over participants.

To investigate the topographical spread of coherence for different image durations, we compared source-level activity during stimulus presentation to baseline activity. We computed the source-level stimulus vs. baseline coherence difference for each of the five image durations. We defined the brain area of interest for each image duration as including all source locations that had an activity value of 50% or higher of the peak activity value for that condition. We also extracted the average y coordinate for each area of interest. This y coordinate was computed by taking the mean of the y coordinates of all source locations belonging to the area for a specific condition.

### Statistical analysis

#### Behavioral results

Mean reaction time and accuracy were first calculated within participant per condition. Then, a two-way repeated measures ANOVA assessed the differences in reaction time, and another two-way repeated measures ANOVA evaluated the differences in accuracy. There were two independent variables, familiarity (two levels: familiar and novel) and image duration (five levels: 50 ms, 100 ms, 150 ms, 200 ms, and 300 ms).

#### Overall amplitude in ERFs

To statistically assess the MEG activity difference between familiar and novel trials and control for multiple comparisons, we applied cluster-based permutation tests (Maris and Oostenveld, 2007), as implemented by FieldTrip (Oostenveld et al., 2011). The tests were carried out on the time period from 0 ms (the onset of the first distractor) and 1200 ms, over all sensors, and with 1000 permutations. For each sensor and time point, the MEG signal was compared univariately between two conditions, using a paired *t* test. Positive and negative clusters were then formed separately by grouping spatially and temporally adjacent data points whose corresponding *p* values were lower than .05 (two-tailed). Cluster-level statistics were calculated by summing the *t* values within a cluster, and a permutation distribution of this cluster-level test statistic was computed. The null hypothesis was rejected if the largest cluster in the considered data was found to be significant, which was the case if the cluster’s *p* value was smaller than .05 as referenced to the permutation distribution.

#### Influence of familiarity and image duration on coherence

To assess how image duration affected the topographical spread of coherence (see above for how this was computed), we conducted a one-way ANOVA with five levels, which were the five image durations. To assess the influence of familiarity and image duration on coherence with the frequency of interest (quantified as source-level coherence as explained above), we performed a two-way repeated measures ANOVA with two factors, familiarity (two levels) and image duration (five levels). Post-hoc *t*-tests were used to assess each pair-wise comparison. Coherence values for each condition were taken from the general visual system region of interest (ROI), which was constructed by taking the union of the five condition-specific areas of entrainment (see Results). Since coherence data are skewed, we log-transformed them to facilitate statistical comparisons between conditions. Results remain qualitatively unaltered if this transformation is not applied.

Next, we computed coherence values for each image duration within ROIs defined separately for each image duration (in contrast to the general visual system ROI described above). We computed the difference in coherence between familiar and novel trials for each image duration in each ROI. Each ROI was based on a stimulus vs. baseline comparison for that image duration, and each successive area did not include the previous one (e.g., the ROI for the 150 ms condition did not include any locations belonging to the 50 or 100 ms conditions). To test how image duration and ROI influenced the magnitude of difference in coherence, we ran a two-way repeated measures ANOVA with two factors, image duration (three levels: 50 ms, 100 ms, and 150 ms) and ROI (five levels). Moreover, we fitted regression lines for each image duration across the five ROIs, and we conducted an analysis of covariance (ANCOVA) to compare the slopes of the regression coefficients for each image duration. For each of these analyses, we did not test the 200 ms and 300 ms conditions because they did not show a significant difference in coherence between familiar and novel trials when tested with the general visual system ROI approach described above (see Results).

#### Correlation between dynamic range and reaction times

To test for the presence of a relationship between coherence (and thus dynamic range) and RT, we computed the correlation between the difference in coherence between familiar and novel trials (familiar-novel) and the difference in RT (novel-familiar). We computed five Pearson’s correlation coefficients, one for each image duration.

#### Influence of familiarity and image duration on target-related amplitude

To determine whether familiarity and image duration influenced ERF amplitude for the target stimulus, we performed a two-way repeated measures ANOVA with two factors, familiarity (two levels) and image duration (five levels) on the time window from 0 ms (target onset) to 300 ms and including all occipital sensors. Post-hoc *t*-tests were conducted to assess each pair-wise comparison. Since combined planar-transformed ERF data are skewed, we log-transformed them to facilitate statistical comparisons between conditions. Results remain qualitatively unaltered if this transformation is not applied.

## Results

We measured MEG activity while participants viewed target stimuli in a stream of familiar or novel distractors. Image duration varied per trial; images were shown for 50, 100, 150, 200, or 300 ms. Participants’ task was to categorize a familiar target image as animate or inanimate as quickly as possible.

### Behavioral Results

#### Behavioral performance improved with distractor familiarity and image duration

Target categorization accuracy was higher when distractors were familiar compared to novel images (*F*_1,36_ = 60.73, *p* = 3.07e-09) and higher for longer image durations (*F*_4,144_ = 130.66, *p* < 1e-15) (see Figure 1B). Moreover, the effect of familiarity was most pronounced for the most challenging rapid visual streams, as indicated by a familiarity by duration interaction (*F*_4,144_ = 24.45, *p* = 1.89e-15).

Target categorization speed was also faster when distractors were familiar compared to novel (*F*_1,36_ = 439.35, *p* < 10e-15) and faster for longer image durations (*F*_4,144_ =166.59, *p* < 10e-15 (see Figure 1B). Again, the effect of familiarity was most pronounced for the most challenging rapid visual streams, as indicated by a familiarity by duration interaction (*F*_4,144_ = 48.27, *p* < 10e-15).

At the end of the MEG session, participants’ knowledge of the distractor image familiarity was assessed. On average, participants correctly identified the familiar images in 91.6% of trials (*SD* = 8.9%), showing that they were clearly aware of the familiarity manipulation.

### MEG Results

#### Novel stimuli led to higher overall activity than familiar ones

In order to investigate the time courses of each trial type, we computed event-related fields (ERFs) (Figure 2A). There was a marked difference in the overall amplitude of the signal, with novel items leading to higher activity than familiar ones (*p* < 1e-03 for all image durations), as found previously in Manahova et al. (2018). The power spectra for the evoked responses and corresponding topographies for each condition are illustrated in Figure 2B. The phase-locked power at the driving frequency approximates the dynamic range of the response (Manahova et al., 2018).

**Figure 2.**
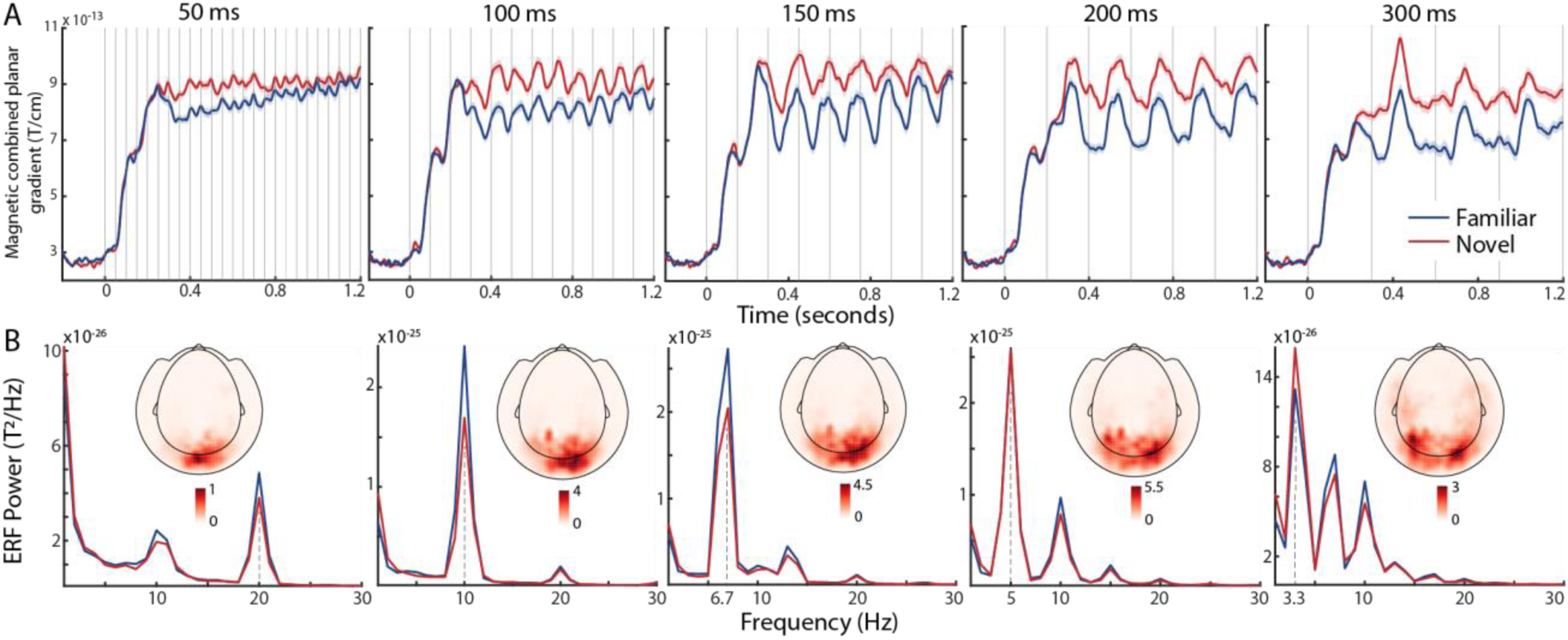
Familiarity effect in sensor-level data. (A) MEG activity (magnetic fields with planar combination) over time for familiar (blue) and novel (red) conditions, separately for each image duration (50, 100, 150, 200, and 300 ms). Activity was averaged over all occipital sensors. Shaded areas are error bars illustrating within-subject SEM for familiar (light blue) and novel (light red) conditions. Vertical lines denote the onset of each image. (B) Power spectra for the event-related fields (ERFs) for familiar (blue) and novel (red) conditions, separately for each image duration (50, 100, 150, 200, and 300 ms). Dotted vertical lines denote the frequency of interest for each image duration. Activity was averaged over all occipital sensors. Topography plots show the spatial distribution of the mean activity of familiar and novel trials for the corresponding image duration. The color bars of the topography plots are of the same units as the y axis of the power spectra.

#### Anatomical location of entrainment

Next, we aimed to identify the anatomical location of entrainment. As a measure of entrainment (i.e., phase-locked power at the stimulus frequency), we computed the coherence between the signal for a specific image duration and a synthetic reference signal (i.e., a cosine function) at the stimulus frequency. We subsequently contrasted the resulting coherence values between stimulus and baseline periods. The resulting five areas of entrainment are depicted in Figure 3A.

**Figure 3.**
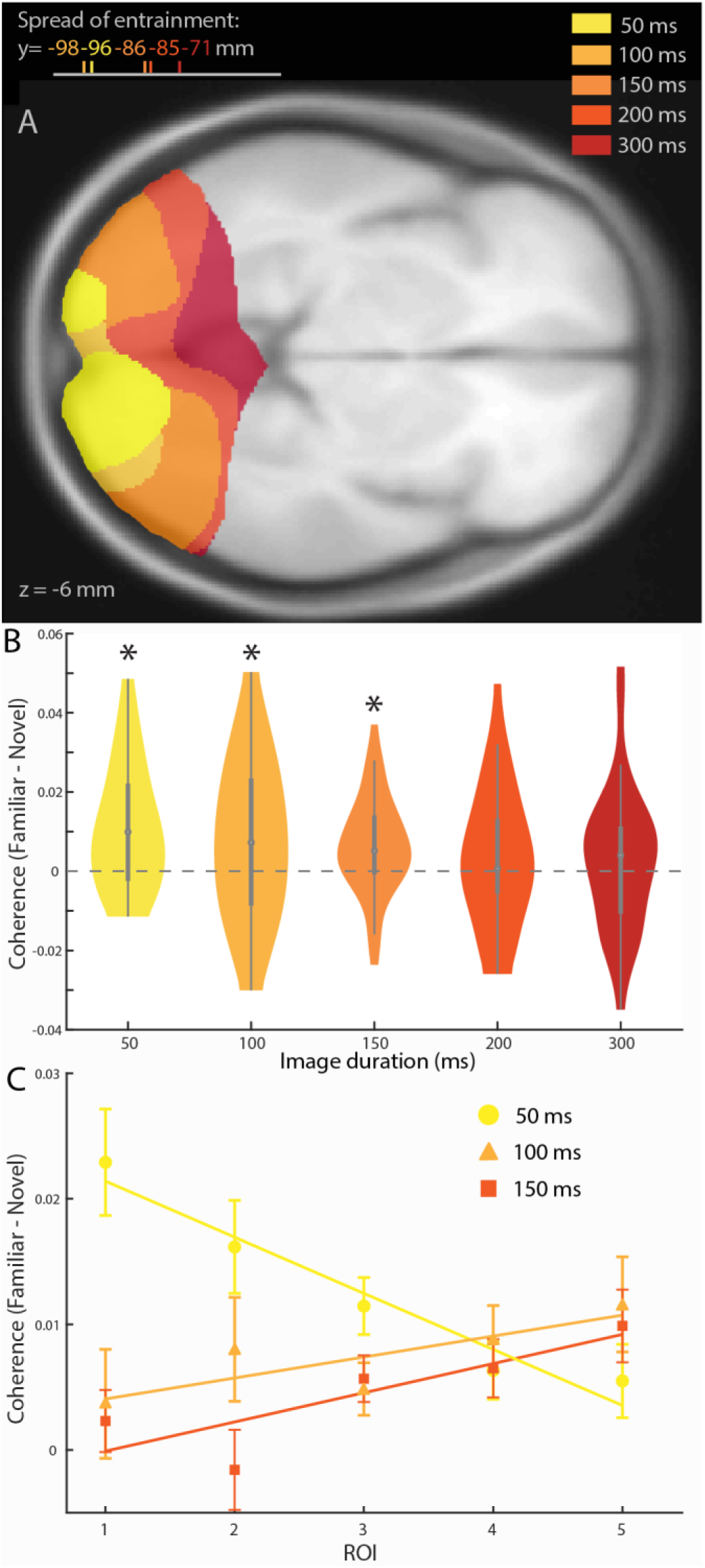
(A) Anatomical location of entrainment in the brain for activity resulting from source reconstruction for stimulus vs. baseline comparison. The anatomical location of activity was estimated separately for each image duration (50, 100, 150, 200, and 300 ms) and, for visual representation, thresholded at 50% of peak value for each condition. The average y coordinates for each image duration are shown in the top left corner. (B) Difference in coherence between familiar and novel trials, separately per image duration. Stars denote significance level at alpha = 0.05. Dashed horizontal line shows the level of no difference. (C) Difference in coherence between familiar and novel trials for different image durations per ROI. Each ROI is based on the stimulus vs. baseline comparison for the corresponding image duration; i.e., ROI 1 corresponds to the 50 ms condition, ROI 2 corresponds to the 100 ms condition, etc. An ROI does not contain any of the locations belonging to the previous ROI; e.g., ROI 3 does not contain any locations belonging to ROIs 1 or 2. Error bars denote within-subject SEM. Lines show regression fits for each image duration.

Strong stimulus entrainment was observed within the ventral visual stream. The topography of entrainment generally spread more anteriorly as the image duration became longer. Each successive area of entrainment (i.e., each area corresponding to a longer image duration) peaked in a progressively more downstream region of the visual hierarchy. Indeed, the average y coordinate shifted more anteriorly as the image duration lengthened (*F*_4,184_ = 21.02, *p* = 3.08e-14), as shown by a one-way ANOVA. Note that this is also in line with the sensor-level topographies of phase-locked power (Figure 2B).

#### Familiar images led to higher dynamic range than novel ones

Next, we aimed to determine whether familiar and novel stimuli lead to significantly different coherence, and thus dynamic range, at each stimulus frequency. We constrained this analysis to a region of interest corresponding to the visual system as identified by our data. Namely, we combined the five areas of entrainment shown in Figure 3A into one region of interest (ROI).

First, we focused on the coherence averaged over this single ROI encompassing a large part of visual cortex. Familiar distractors were associated with stronger coherence than novel ones (*F*_1,36_ = 22.09, *p* = 3.75e-05), and coherence also differed as a function of image duration (*F*_4,144_ = 7.07, *p* = 3.16e-05), but the interaction was not significant (*F*_4,144_ = 1.10, *p* = 0.36) (see Figure 3B). We found that coherence was significantly higher for familiar than for novel trials when images were presented for 50 ms (*t*_36_ = 4.23, *p* = 1.50e-04), 100 ms (*t*_36_ = 2.39, *p* = 0.02), and 150 ms (*t*_36_ = 3.08, *p* = 0.004). There was no significant difference between familiar and novel for images lasting for 200 ms (*t*_36_ = 1.14, *p* = 0.26) or 300 ms (*t*_36_ = 0.89, *p* = 0.38). Thus, the dynamic range was higher for familiar than novel stimuli. The data suggest that this difference was particularly prominent for the short and medium image durations (50, 100, and 150 ms), although the lack of a significant interaction indicates there is no statistical support for significantly stronger entrainment for short compared to long (200 and 300 ms) image durations.

#### Familiarity-induced truncation shifted topographically depending on image duration

Next, we shifted our attention from the single visual system ROI to individual ROIs defined for each image duration. To assess how the topographical distribution of familiarity-induced truncation changed with image duration, we quantified the difference in coherence between familiar and novel trials in the five regions of interest, each based on the area of entrainment for an image duration (see Figure 3C). Importantly, each successive area did not include the previous one (e.g., the ROI for the 150 ms condition did not include any locations belonging to the 50 or 100 ms conditions). For this analysis, we examined activity for the 50, 100, and 150 ms conditions because they showed significant differences in coherence between novel and familiar distractors when tested within the general visual system ROI (see Figure 3B). The different conditions exhibited different patterns of signal truncation across ROIs, as evidenced by a significant interaction (*F*_4,144_ = 4.17, *p* = 9.83e-05). To understand this interaction effect better, we conducted an analysis of covariance (ANCOVA), fitting regression lines across ROIs for each image duration (see Figure 3C) and then comparing the regression coefficients to each other. We found that signal truncation decreased significantly with anteriority for the 50 ms condition (*t* = −4.24, *p* < 0.01), while we observed a trend in the opposite direction for the 100 ms condition (positive slope but not significantly different from zero; *t* = 1.80, *p* = 0.07). For the 150 ms condition, signal truncation increased significantly with anteriority (*t* = 2.44, *p* = 0.02). Thus, for the 50 ms condition, signal truncation peaked in the earliest ROI and decreased in successive ones, while for the 100 and 150 ms conditions, signal truncation was low in early ROIs and increased in successive ones.

#### No relationship between dynamic range and reaction times

We found a clear reaction time benefit for familiar stimuli compared to novel ones, as well as a clear difference in signal truncation. This raises the question of whether these two effects are correlated across participants. To quantify this, we computed the correlation between the difference in coherence on familiar and novel trials (familiar-novel) and the difference in RT on familiar and novel trials (novel-familiar) across subjects. We computed five correlation coefficients, one for each image duration. None of these correlations were significant (all *ps* > 0.1).

#### Stronger processing of target image when embedded in familiar distractors

We assessed how strongly a target was processed when it was embedded in familiar or novel distractors by examining the overall amplitude of target-evoked activity in the time window from 0 ms (target onset) until 300 ms over occipital sensors, as seen in the target-related ERFs (see Figure 4). The target was processed more strongly when it was embedded in familiar distractors rather than novel ones (*F*_1,36_ = 233.72, *p* < 10e-15). While this effect was not equally strong for all image durations (familiarity X duration interaction: *F*_4,144_ = 5.27, *p* = 5.46e-04), post-hoc tests confirmed that it was robustly present for all image durations (50 ms: *t*_36_ = 11.46, *p* = 1.43e-13; 100 ms: *t*_36_ = 10.08, *p* = 4.99e-12; 150 ms: *t*_36_ = 11.32, *p* = 2.04e-13; 200 ms: *t*_36_ = 8.14, *p* = 1.11e-09; 300 ms: *t*_36_ = 5.78, *p* = 1.35e-06). Therefore, the target stimulus resulted in stronger evoked responses for familiar than for novel trials, regardless of image duration.

**Figure 4.**
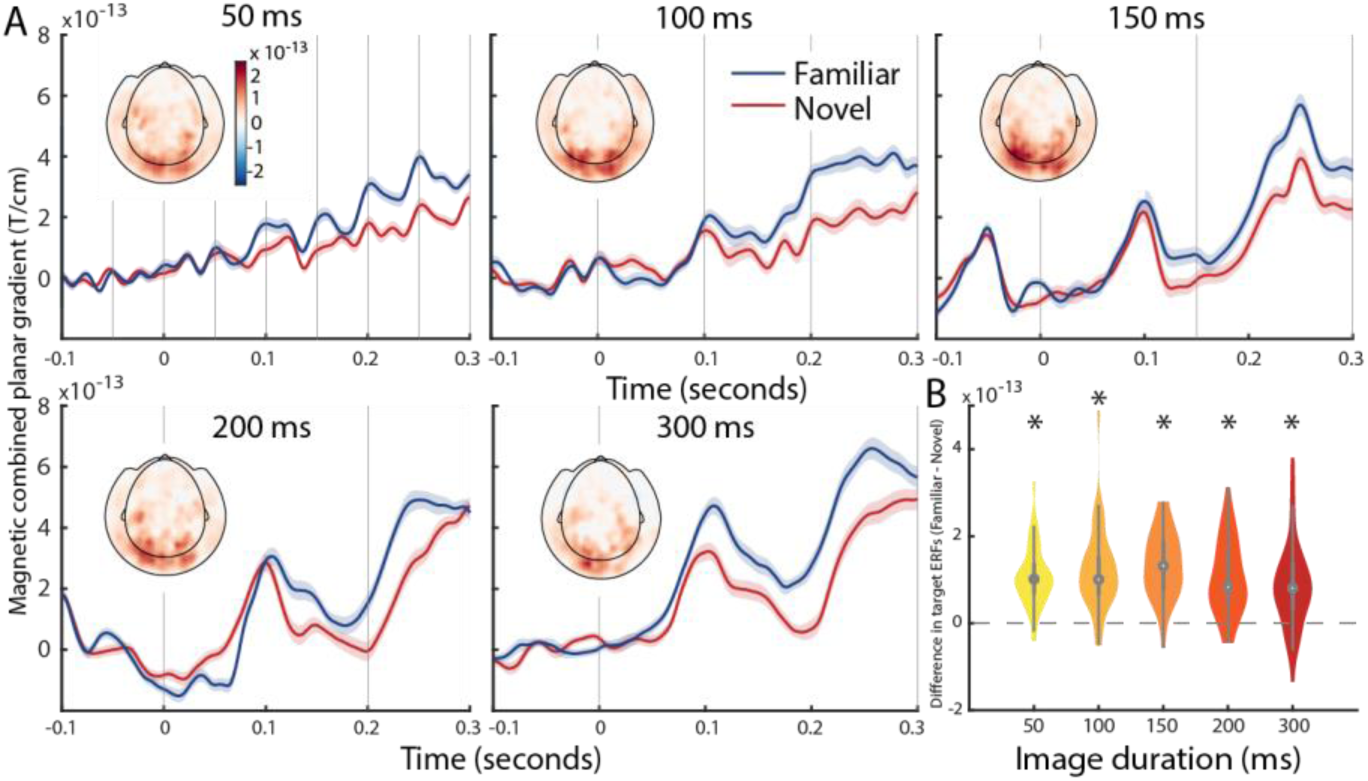
Target-related activity in sensor-level data. (A) MEG activity (combined planar gradient) over time for familiar (blue) and novel (red) conditions, separately for each image duration (50, 100, 150, 200, and 300 ms). Activity was averaged over all occipital sensors. Shaded areas are error bars illustrating within-subject SEM for familiar (light blue) and novel (light red) conditions. Vertical lines denote the onset of each image. Topography plots show the spatial distribution of the difference in target-related activity between Familiar and Novel streams (Familiar – Novel) separately for each image duration. Activity was averaged over the time window from 0 ms (target onset) until 300 ms. The color bars of the topography plots are of the same units as the y axis of the ERFs, and the scale of the color bars is the same for all topography plots. (B) Difference in target-related activity between familiar and novel trials (Familiar – Novel), separately per image duration. Stars denote significance level at alpha = 0.05. Dashed horizontal line shows the level of no difference.

## Discussion

In this study, we investigated the temporal and spatial dynamics of signal truncation and whether this phenomenon bears relationship to participants’ ability to detect target items within a stream of visual distractors. In short, we found truncation of neural activity for familiar input to be the strongest when image streams were presented rapidly (images last between 50 and 150 ms each). For the shortest image duration (50 ms), signal truncation was localized to early visual cortex, while successively longer image durations were linked to progressively later visual areas. Furthermore, the neural processing of the target stimulus was stronger when targets were embedded in familiar streams compared to novel ones. Behavioral performance was also better for targets in familiar streams, especially for short image durations.

A wealth of research has demonstrated that familiarity affects neural processing, the most commonly observed effect being familiarity suppression (Fahy et al., 1993; Li et al., 1993; Sobotka and Ringo, 1993; Xiang and Brown, 1998; Freedman et al., 2006; Mruczek and Sheinberg, 2007; Anderson et al., 2008; Woloszyn and Sheinberg, 2012; Huang et al., 2018). Viewing an object image repeatedly and becoming familiar with it results in reduced spiking activity in inferotemporal cortex (IT) in monkeys (Miller et al., 1991) and reduced hemodynamic activity in human LOC as measured with fMRI (Grill-Spector et al., 2006). This reduction of the population response may indicate sharper neuronal tuning and a sparser population representation for familiar than for novel images (Freedman et al., 2006; Woloszyn and Sheinberg, 2012). Our data are in accordance with this reduction of activity for familiar stimuli: we found a sustained higher amplitude response for novel than for familiar items in the event-related fields (ERFs) (see Figure 2A).

Image familiarity does not only influence the magnitude of the neural response, but also truncates the neural response (Meyer et al., 2014), such that neural activity returns to baseline levels more quickly for a familiar than a novel image. This puts neurons in a state of readiness to respond to new input more rapidly. We observed signal truncation throughout the ventral visual system (Figure 3A). Interestingly, the topographical distribution of the effect shifted anteriorly and laterally as image duration increased. Perhaps surprisingly, signal truncation was robustly present in early visual cortex, particularly for very rapid streams (50 ms per image). In line with this, it has recently been observed that correlates of visual familiarity are observed as early as macaque area V2 (Huang et al., 2018).

Since signal truncation of distractors may benefit target processing, we explored whether signal truncation was related to target processing and behavioral performance. Indeed, we found that while novel distractors showed a higher amplitude than familiar distractors (Figure 2A), target-related activity was higher when the targets were embedded in familiar images than in streams of novel images (Figure 4). In terms of behavioral performance, participants were markedly faster and more accurate in categorizing the target when the target was embedded within familiar distractors, suggesting a direct relationship between signal truncation of the distractors and behavioral performance. However, a between-subject correlation analysis showed no reliable correlation between truncation and behavior. Nevertheless, both the stronger evoked responses and the enhanced behavioral performance for targets embedded in familiar streams suggests a benefit for the processing of visual input, possibly stemming from the signal truncation of the distractors. This is in accordance with findings showing that familiar distractors are less salient (Jutras and Buffalo, 2010; Ghazizadeh et al., 2016) and less disruptive than novel ones (Mruczek and Sheinberg, 2007).

We hypothesized that signal truncation would be prevalent in early visual cortex for short image durations and occur in later visual areas as images are shown for longer. We based this idea on the fact that early regions reach their peak activity relatively soon after stimulus onset, while more downstream regions show peak spiking later (Dinse and Krüger, 1994; Nowak and Bullier, 1997), and that higher-order cortical regions have a slower intrinsic timescale than early sensory regions (Murray et al., 2014). Moreover, regions lower in the visual hierarchy accumulate input for shorter image durations than downstream areas (Hasson et al., 2008; Honey et al., 2012). Our results are in accordance with these notions. First, we found overall entrainment (from a stimulus vs. baseline contrast) that showed an anterior and lateral topographical shift with increasing image duration. Second, the spatial localization of the difference in signal truncation between familiar and novel stimuli also shifted to more downstream areas as images were shown for longer, with the 50 ms condition showing the strongest signal truncation difference for early ROIs, while the 100 and 150 ms conditions demonstrated stronger signal truncation difference for more downstream ROIs. Therefore, both overall entrainment and the signal truncation difference showed a topographical distribution that shifted up the brain’s visual processing hierarchy along with increasing image duration.

It is intriguing to speculate how signal truncation can be observed for such short image durations as 50 ms and in early visual cortex. This implies that familiarity may be encoded in early visual cortex somehow, which may sound implausible since familiarity suppression effects are commonly established in IT (e.g., Li et al., 1993). However, Huang et al. (2018) showed that a type of familiarity suppression is also found in macaque V2, which suggests that early visual cortex may encode familiarity. Alternatively, the signal truncation effect we observe in early visual areas may be the result of feedback from higher-order areas. In our current analysis, and in accordance with previous work (Manahova et al., 2018), we quantify signal truncation by computing coherence over a 1000 ms time window, so we are unable to detect at which time points in the image stream truncation is present. It is possible, however, that the visual evoked response may be truncated only after the first few stimuli in a stream are presented, which may indicate the involvement of higher areas. After all, the processing of an image continues after other images are presented even when images are presented as briefly as 17 ms (Mohsenzadeh et al., 2018), suggesting an effect such as signal truncation may build up in magnitude as a challenging visual stream is being processed.

In the current study, image durations of 200 ms and 300 ms did not elicit a significant difference in signal truncation. Future research could use more complex or naturalistic (Kayser et al., 2004) images because such stimuli, which require more extensive and demanding processing, may engage regions higher up in the visual hierarchy (Murray et al., 2014), thus leading to signal truncation differences even if images are shown for 200 ms or longer.

Although we compared familiar and novel images, our familiarity manipulation involved both familiarity and recency. Xiang and Brown (1998) define familiarity as whether an object was seen during a previous session, usually on a previous day, and recency as whether an object has been seen previously during the same recording session. While these effects are clearly related, as they refer to whether the system has been exposed to this stimulus before, they denote different time scales (Fahy et al., 1993) and may rely on different mechanisms. Familiarity, related to a longer time scale, requires plasticity changes in the neural network that encodes the stimulus. Recency, related to a shorter time scale, is more similar to repetition suppression or adaptation effects and may result from synaptic depression or adapted input from other neurons in the same network (Vogels, 2016). In the current study, we manipulated both familiarity and recency in our familiar condition, meaning that we are unable to distinguish the effects of these two phenomena. It would be interesting to further disentangle the respective contributions of recency and familiarity to signal truncation.

In conclusion, familiarity with a visual stimulus leads to a truncation of neural activity in the visual system, and this truncation was strongest for the fastest presentation speeds. Moreover, this truncation for familiar distractor stimuli is associated with stronger target processing and behavioral improvements in perceptual categorization, suggesting a functional role of this phenomenon.

## Acknowledgements

This work was supported by The Netherlands Organisation for Scientific Research (NWO Vidi grant 452-13-016 awarded to FPdL; NWO Veni grant 016.Veni.198.065 awarded to ES; NWO Research Talent grant 406-16-525 awarded to MEM) and the EC Horizon 2020 Program (ERC starting grant 678286, ‘Contextvision’ awarded to FPdL). The authors thank Louise Barne and Ashley Lewis for helpful comments on this manuscript.

